# Stress-induced dendritic branching in *C. elegans* requires both common arborization effectors and stress-responsive molecular pathways

**DOI:** 10.1101/808337

**Authors:** Rebecca J. Androwski, Nadeem Asad, Janet G. Wood, Allison Hofer, Steven Locke, Cassandra M. Smith, Becky Rose, Nathan E. Schroeder

## Abstract

Stress influences the shape of dendritic arbors in neurons. During the stress-induced dauer stage of *Caenorhabditis elegans*, the IL2 neurons arborize to cover the anterior body wall. In contrast, the FLP neurons arborize to cover the anterior body wall during non-dauer development. Previous work showed that the membrane-bound receptor DMA-1 regulates FLP branching as part of a larger protein complex. Using forward genetics, we show that the IL2 neurons also use the DMA-1 complex to regulate branching. To understand the coordination of the IL2s and FLPs we conducted a time-course examination of FLPs and found previously undescribed branching patterns indicating a neighborhood effect wherein the FLPs and IL2s in the anterior have differential branching compared to the more posteriorly located PVD arborizing neurons. To determine how the IL2s and FLPs differentially regulate branching, we examined several regulators of DMA-1 localization. We show that the unfolded protein response sensor IRE-1, required for FLP branching, is only required for dauer-specific branching at elevated temperatures. Interestingly, we found that *ire-1* mutants have broad, organism-wide temperature-dependent effects on dauer remodeling, suggesting a previously undescribed role for IRE-1 in phenotypic plasticity. We also found that defects in other regulators of dauer remodeling including DAF-16/FOXO, DAF-9/Cytochrome P450, and DAF-18/PTEN are required for proper IL2 arborization, but dispensable for FLP branching. Interestingly, we find that TOR adaptor protein DAF-15/RAPTOR is both required for promoting IL2 branching and inhibiting precocious development of the FLPs. Our results demonstrate specific genotypic by environmental interactions regulating dendrite arborization.

**SIGNIFICANCE STATEMENT:** Neurons have extensions called dendrites that receive information. Dendrites are often elaborately shaped with many branches. Adverse stress can reduce branching in some neurons, while increasing it in others. How stress can cause some neurons to change shape is unclear. We previously found a set of neurons in the head of the well-studied roundworm *C. elegans* that undergo reversible branching following exposure to specific adverse environmental conditions. Using various genetic tools, we find that branching in these neurons is controlled by a combination of branching genes common to many neuron types and others that only regulate branching in stress-responsive neurons. Our data demonstrate how experiencing stress acts through genetics pathways to cause changes to specific neurons.

## INTRODUCTION

Dendrite morphology is crucial for efficient neural signaling. Recent years have seen increased attention given to the molecular basis of dendritic arborization using both cell-culture and model organisms. Experience-driven remodeling of a dendrite can contribute to both the size and complexity of dendritic arbors in a diverse range of species. Depending on neuron class these changes can include both increasing and decreasing dendritic volume (1–3). For example, dendrites in the basal lateral amygdala undergo hypertrophic growth following chronic stress, whereas those of the hippocampus atrophy. However, little is known about the molecular mechanisms that cause differential arborization in different neuron classes during normal physiological conditions versus stress-exposed conditions. Similarly, what, if any, coordination occurs among distinct neuron types is unknown.

The nematode C*aenorhabditis elegans* is one of several leading models for studying dendritic arborization. Its transparent body, well-described nervous system, and abundant genetic tools available for its study comprise a robust system for understanding the molecular basis of neuron morphology. Under standard well-fed laboratory conditions, *C. elegans* develops from an embryo to adult within three days (4). During development, the PVD and FLP neurons undergo progressive arborization of the midbody and head, respectively (Fig. S1). The PVDs and the FLPs reach their maximum outgrowth during the fourth larval stage (L4) immediately before entering reproductive maturity (5).

Under conditions of reduced food availability and high population density, *C. elegans* can enter into an alternative juvenile stage called dauer (6). Dauers undergo extensive remodeling of various tissues leading to a non-feeding stress-resistant stage. We previously demonstrated that the inner-labial (IL2) neurons undergo dendrite arborization of the head during dauer formation (3). During non-dauer stages, the IL2s are simple bipolar neurons with a single unbranched dendrite (7). During dauer formation, the four subdorsal and subventral IL2s undergo extensive branching (Fig. S1). When returned to favorable environmental conditions, dauers recover to resume the standard developmental course towards adulthood. During dauer recovery, the IL2 arbors are resorbed and return to their non-dauer morphology.

We previously demonstrated that the proprotein convertase KPC-1 is required for both dauer-specific IL2 and PVD/FLP arborization (3). Subsequent work on the PVDs demonstrated an interaction between KPC-1 and DMA-1, which encodes a membrane-bound leucine-rich repeat receptor (8). DMA-1 acts as part of a protein complex with the hypodermally-localized L1CAM homolog SAX-7 and MNR-1, a conserved transmembrane protein (9, 10). Concurrently, the diffusible LECT-2, homologous to leukocyte cell-derived chemotaxin 2, is secreted from surrounding muscle to increase the binding affinity of the DMA-1 complex (11). The DMA-1 complex in the PVDs and FLPs is regulated through diverse cell functions such as the unfolded protein response and membrane trafficking (12–15).

Here, we use both forward and reverse genetics to demonstrate that dauer-specific IL2 arborization uses shared and distinct molecular mechanisms from the PVDs and FLPs to control arborization. Specifically, we find that while the PVDs, FLPs, and IL2s all use the DMA-1 complex, the regulation of this complex differs between the PVD/FLPs and IL2s. Given the shared location of the FLPs and IL2s in the head of *C. elegans*, we examined the structure of the FLPs throughout development and observed architectural differences between the anterior FLPs and IL2s and the posterior PVDs, emphasizing the importance of neighborhood in determining neuronal structure. Finally, we uncover a temperature-mediated regulation for the unfolded protein response sensor IRE-1 in IL2 branching and describe a collection of new and previously described dauer-remodeling genes that control arborization in the IL2s.

## RESULTS

### The DMA-1 complex is required for dauer-specific IL2 arborization

In order to determine mechanisms controlling dauer-specific IL2 arborization, we performed a mutagenesis screen. From this screen, we identified a recessive mutation *cbc1*, which leads to disorganized and truncated IL2 branching during dauer with complete penetrance (Fig. 1A). Using traditional genetic mapping, we placed *cbc1* near *dma-1*, a leucine-rich receptor encoding protein required for branching in the PVD neurons. We found that *cbc1* fails to complement the deletion mutant *dma-1(tm5159)* for IL2 branching. Subsequent sequencing of the *dma-1(cbc1)* locus identified a single T to A missense mutation resulting in a leucine to histidine change within the leucine-rich repeat of DMA-1 (Fig. 1B). We found no obvious difference in the severity of branching defects between *dma-1(cbc1)* and the deletion allele *dma-1(tm5159)*, suggesting that *dma-1(cbc1)* is a strong loss-of-function allele (Fig. S2A).

**Figure 1.**
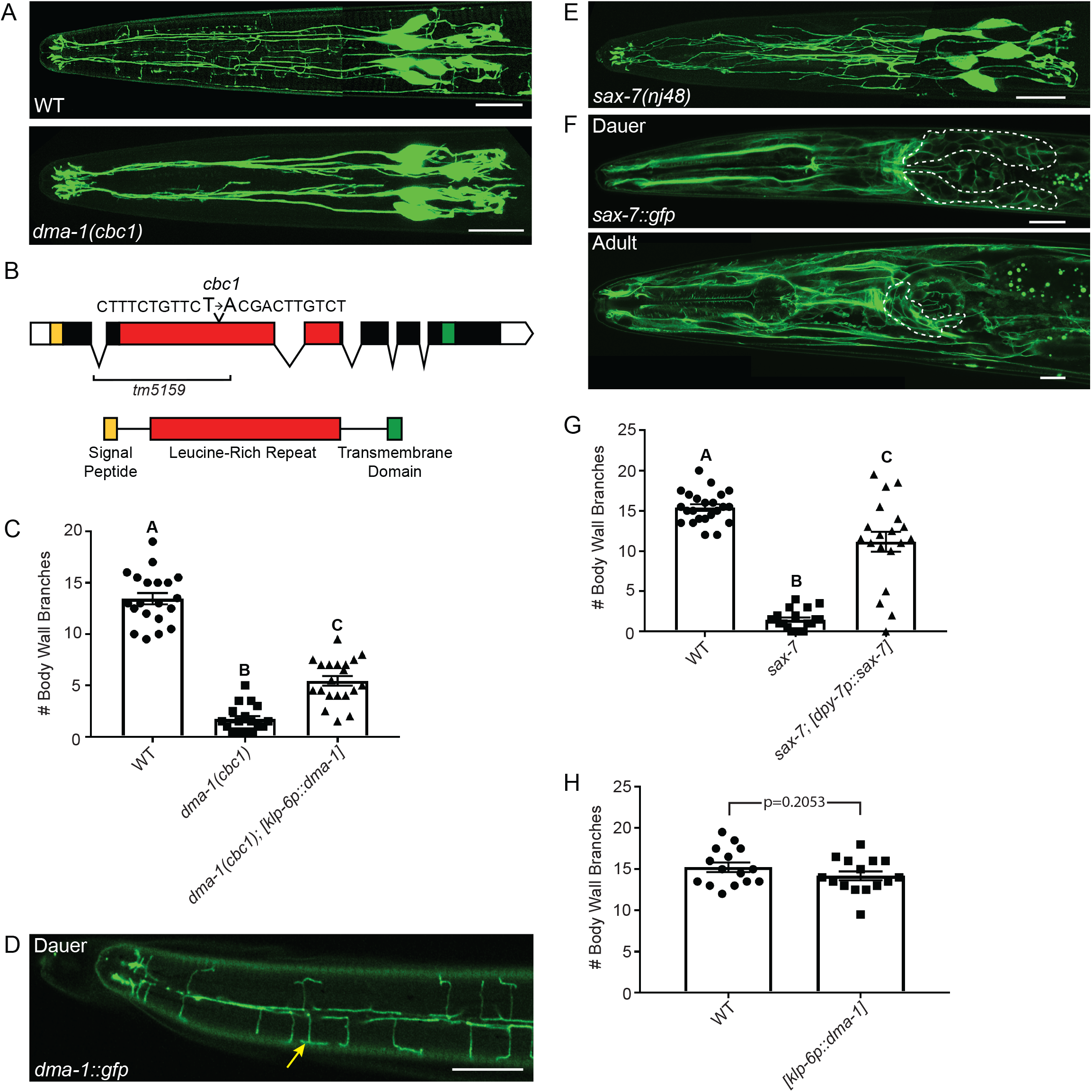
The DMA-1 complex is required for IL2 arborization. (A) Z-projection confocal micrographs of dauers expressing expressing gfp in the IL2 neurons. The IL2 neurons arborize extensively in wild-type dauers, forming body wall branches that cover each muscle quadrant. *dma-1(cbc1)* mutants have severe defects in IL2 arborization. (B) Gene schematic of *dma-1. dma-1(cbc1)* is a point mutation that results in a T to A transversion within the leucine-rich repeat (red) region of *dma-1*. The previously described deletion allele *dma-1(tm5159)* removes much of the LRR domain (9). (C) Quantification of the number of IL2 branches along the body wall during dauer in wild-type (n=19), *dma-1*(n=18), and *dma-1; klp-6p∷dma-1* (n=20). Expression of *dma-1* in the IL2 neurons using the *klp-6* promoter rescues the *dma-1* mutant branching phenotype. (D) Z-projection confocal micrograph of dauer expressing *dma-1∷gfp* expression under the *dma-1* promoter. *dma-1∷gfp* localizes to the IL2 dendrite (yellow arrow) during dauer. This substack of images includes the dendrites located at the body wall. (E) Z-projection confocal micrograph of *sax-7(nj48)* mutant dauer expressing gfp in the IL2 neurons. *sax-7(nj48)* mutants have severe defects in IL2 arborization. (F) Confocal micrograph of *sax-7∷gfp* expression under its endogenous promoter during dauer and adult. A dorsal-ventral view of a dauer (top) from a single z-plane shows widespread expression of *sax-7∷gfp* in the cell membranes of neuronal (anterior ganglia indicated by dashed line) and non-neuronal tissues. A lateral view of an adult (bottom) shows a similar expression pattern to the dauer. (G) Quantification of the number of IL2 body wall branches in *sax-7* mutants and hypodermal-specific rescue of *sax-7*. WT (n=23), *sax-7* (n=19), and *sax-7; dpy-7p∷sax-7* (n=19). (H) Quantification of *dma-1* overexpression during dauer WT (n=15) and *klp-6p∷dma-1* (n=17). For all rescue experiments an ANOVA followed by Tukey’s test for multiple comparisons was used to determine significance except for (H) where a t-test was used. Different letters above bars indicate statistical significance (p<0.0001). Error bars are the standard error of the mean. IL2 neurons are labeled with *klp-6p∷gfp* (A and E). Scale bars, 10 μm.

DMA-1 acts cell-autonomously in the PVDs to regulate arborization and localizes to the PVD dendrite (16). Similarly, IL2-specific rescue of DMA-1 was sufficient to rescue the dendrite phenotype in *dma-1(cbc1)* dauers (Fig. 1C). Furthermore, we observed *dma-1∷gfp* expression in the IL2s during dauer though not during non-dauer stages (Fig. 1D). Altogether, our results suggest that DMA-1 is acting similarly during PVD/FLP arborization and dauer-specific IL2 arborization.

In the PVDs, DMA-1 acts through a protein complex for proper branching (10, 11, 17, 18). *sax-7* encodes a cell adhesion molecule, homologous to mammalian L1CAM, which acts in the surrounding hypodermis to control PVD arborization (9, 10). Loss of *sax-7* leads to reduced and disorganized PVD arborization (9, 10). Similarly, we found that *sax-7* is required for dauer-specific IL2 arborization (Fig. 1E). Examination of a *sax-7∷gfp* reporter revealed widespread expression in dauers, including neuronal and non-neuronal tissue (Figs. 1F and G) (19). Similar to the PVDs (9, 10), we found that hypodermal-specific expression successfully rescued the IL2 branching phenotype in *sax-7* dauers (Fig. 1G). Similarly, we found that *mnr-1* mutants, which is required for PVD and FLP arborization, are also defective for dauer-specific IL2 branching and that these defects could be rescued through hypodermal-specific expression (Figs. S2A and B). Finally, the muscle-secreted protein LECT-2 interacts with the DMA-1 complex to promote PVD branching (11, 18). Consistent with this, we found that loss of *lect-2* causes defects in dauer-specific IL2 branching that can be rescued through muscle-specific expression (Fig. S2A and C). Together, our data suggest that the DMA-1 complex is a common tool for branching in all highly arborized neurons of *C. elegans*.

Due to the central role of DMA-1 in IL2 arborization, we were interested if DMA-1 expression alone was sufficient to generate new branching phenotypes. Previous work showed that overexpression of DMA-1 in the PVDs leads to supernumerary branching (16). However, we found that expression of an IL2-specific DMA-1 construct in a wild-type background did not produce extra branches during dauer (Fig. 1H). Similarly, we found that overexpression of DMA-1 in the IL2s during adult was not sufficient to induce IL2 branching in non-dauers (n=17). Previous work showed that expression of DMA-1 could induce ectopic branching in the non-arborizing PLM touch-receptor neurons adjacent to the PVDs (16). However, we did not observed ectopic branching in the anterior ALM and AVM touch receptor neurons following expression of DMA-1 in the anterior touch receptor neurons during adult (n=32) or dauer (n=32). These data may suggest that neurons in the anterior are more resistant to ectopic branching and corresponds with previous work showing that expression of DMA-1 in the unbranched OLL anterior sensory neurons did not result in increased complexity (16).

### The PVDs, FLPs and IL2s have distinct branching architectures

The neighborhood where a neuron is located can lead to differences in branching (20). As the FLPs share a similar receptive field as the IL2s, we wanted to examine the coordination of branching between these two cell types. The lack of an FLP-specific reporter and the obvious stereotypical morphology of the PVDs has meant that the FLPs have received less attention than PVD arborization. We therefore used Airyscan super-resolution microscopy to examine the architecture of FLP neurons. We found that the FLPs, PVDs and IL2s have distinct dendritic architectures compared to each other. Three major features distinguish the FLPs from the PVDs. First, unlike the regular menorah-like structures of the PVDs, we find that the adult FLPs give rise to opposing menorahs in each body wall quadrant where the “candles” traverse the body wall from both the lateral and dorso-ventral directions (Fig. 2A). This branching pattern is more similar to the IL2 arbor than the PVDs. Second, as a result of several branch points before the FLP dendrites reach the body wall the FLPs can have up to seven orders of branching compared with the four or five found in the PVDs. Lastly, the FLP primary dendrite is already branched in L1 unlike the PVDs, which produce secondary dendrites in L3 (Fig. S3) (21). A detailed description of the adult FLP branching route is given in the figure legend (Fig. 2 and S3).

**Figure 2.**
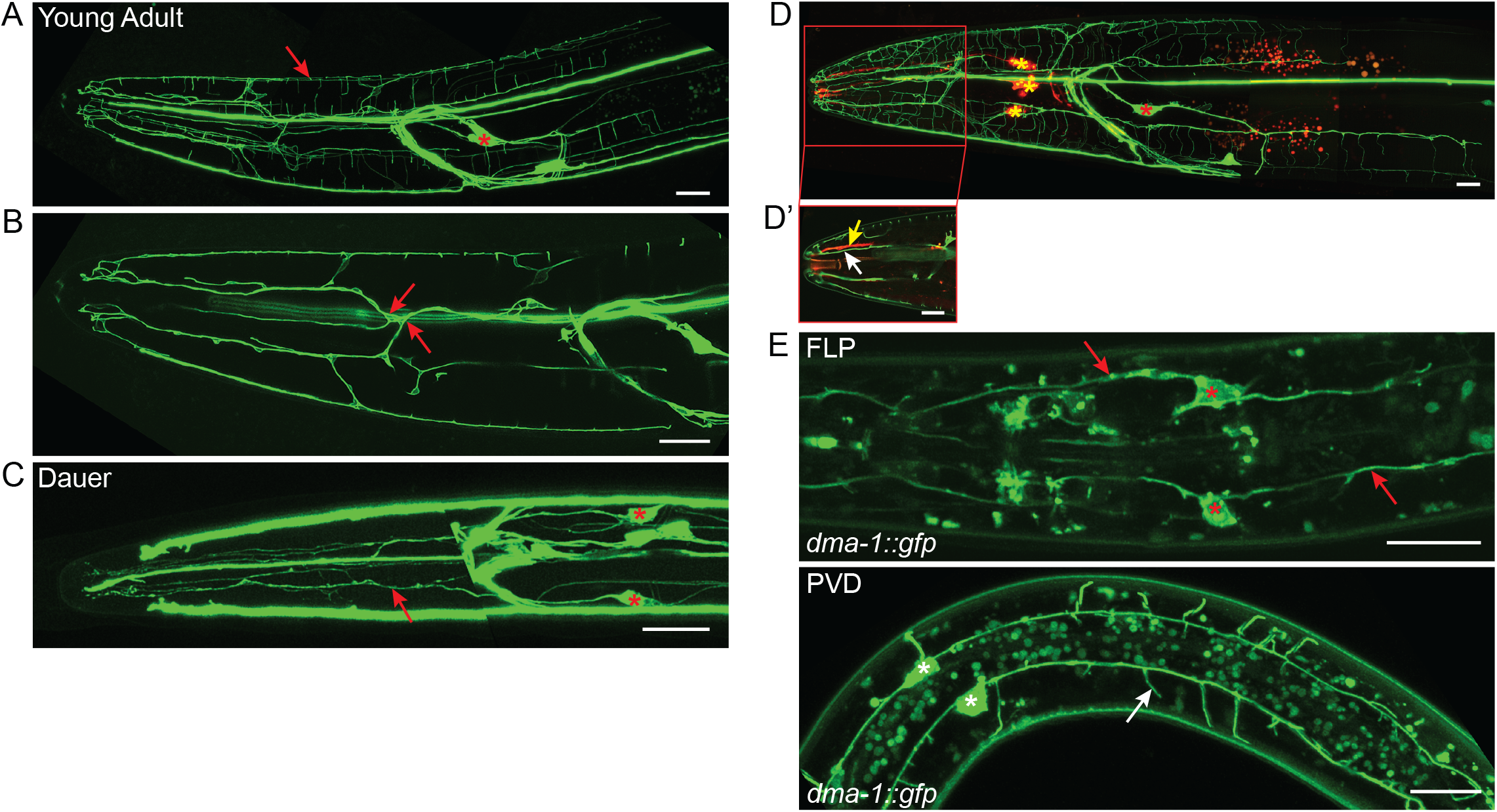
FLP branching architecture. (A-C) Z-projection confocal micrographs of FLP neurons. (A) Lateral view of the adult FLP arbors showing extensive dendritic arborization covering the head of the nematode. The primary dendrite travels along the lateral sensory neuron fascile until anterior of the metacorpus. There it branches and sends secondary processes to the subdorsal and subventral sensory neuron fascicles. A thin primary dendrite continues within the lateral sensory fascicle, but does not reach the nose.The secondary dendrites in the subventral and subdorsal fascicles branch again to send processes to the dorsal or ventral midlines and sublateral lines along the body wall where they will branch to form the body wall candles of the arbor. Red asterisks indicate the FLP cell body, red arrows indicate the FLP dendrite. (B) A subset of slices from a stack of a young adult showing branching of the FLP primary dendrite. Red arrows indicate points where the primary dendrite divides to form secondary dendrites within the subdorsal and subventral sensory neuron fascicles. This branch point exists in L1 (see Fig. S3A) (C) Dorsal-ventral view of the dauer FLP arbors showing minimal branching compared to the adult. Dauers have no FLP body wall branches. Red asterisks indicate the FLP cell body red arrows indicate the FLP dendrite. (D) Co-labeling of IL2 neurons (red) and the FLP neurons (green). Yellow asterisks indicate IL2 cell bodies while red asterisks indicate the FLP cell bodies. (D’) A single image slice showing the proximal relationship between the FLP dendrite (white arrow) and IL2 dendrite (yellow arrow) in the subdorsal sensory neuron fascicle. The FLPs are labeled with with *mec-7p∷gfp*, which also labels with ALM and AVM touch receptor neurons. The IL2 are labeled with *klp-6p∷tdtomato.* (E) Z-projection confocal micrograph of *dma-1∷gfp* expression under its endogenous promoter during dauer. These image substacks include the area surrounding the FLP (top) and PVD (bottom) cell bodies respectively. During dauer, *dma-1∷gfp* localizes to the FLP cell body (red asterisks) and dendrite (red arrows) in the head and the PVD cell body (white asterisks) and dendrites (white arrow) in the body. Scale bars, 10 μm.

The FLPs, which are born embryonically, have already established their primary dendrite prior to hatching (7). However, we find that unlike the PVD and IL2 unbranched primary dendrite, the FLP primary dendrite has a single branch anterior of the metacorpus during early development (L1-L3) forming two parallel, longitudinal processes that likely travel in the subdorsal and subventral sensory neuron fascicles (Fig. 2B). Similar to previous descriptions, we found little additional growth arborization until L4 when rapid growth of both the FLPs and PVDs commence (Fig. S3) (22). As the FLPs share a receptive field with the IL2s, we were particularly interested in the structure of FLPs during dauer. Although several neurons have dauer-specific morphology, we found that the FLPs do not (Fig. 2C) (3, 23, 24). The FLP dendrite retains its L2 morphology during dauer. A similar finding was recently described in the posterior PVD neurons during dauer (25). Similarly, while the non-dauer IL2 dendrites fasciculate with the lower order FLP longitudinal dendrites, the body wall branches of IL2 dauers and FLP non-dauers are strictly separated by development stage (Fig. 2D). Our analysis of FLP and IL2 branching may suggest both neighborhood effects, which create similarities in branch patterns, as well as developmental coordination between the two neuron classes.

### Regulatory mechanisms controlling DMA-1 localization differ between IL2 and PVD/FLP arbors

To understand how the FLPs and IL2s coordinate their branching, we hypothesized that *dma-1∷gfp* would not be expressed in the FLPs during dauer. To our surprise, we found *dma-1∷gfp* prominently expressed in the FLPs and PVDs during dauer (Fig 2E). This result suggests there are additional factors inhibiting FLP and PVD branching in dauers. We, therefore, examined known regulators of the DMA-1 complex. Exocytosis of DMA-1 from the endoplasmic reticulum to the PVD dendritic membrane uses the RAB-like GTPase, RAB-10 and an ortholog to human exocyst complex component 8, EXOC-8. Loss of *rab-10* and *exoc-8* lead to reduced PVD arborization (14, 15). Interestingly, we found that *rab-10* and *exoc-8* are dispensable for dauer-specific IL2 arborization (Fig. 3A). We hypothesized that a RAB-10 paralog could function in the IL2s to transport DMA-1 to the membrane during dauer. Using a phylogenetic examination of human Rab GTPases as a guide (26), we examined mutants of the *rab-10* paralogs *rab-8* and *rab-35* for effects on IL2 arborization. We found no effect from the loss of *rab-8* or *rab-35* on IL2 arborization (Fig. S4A). Our results align with recent findings regarding a reduction in endosome production in the dauer PVDs (25).

**Figure 3.**
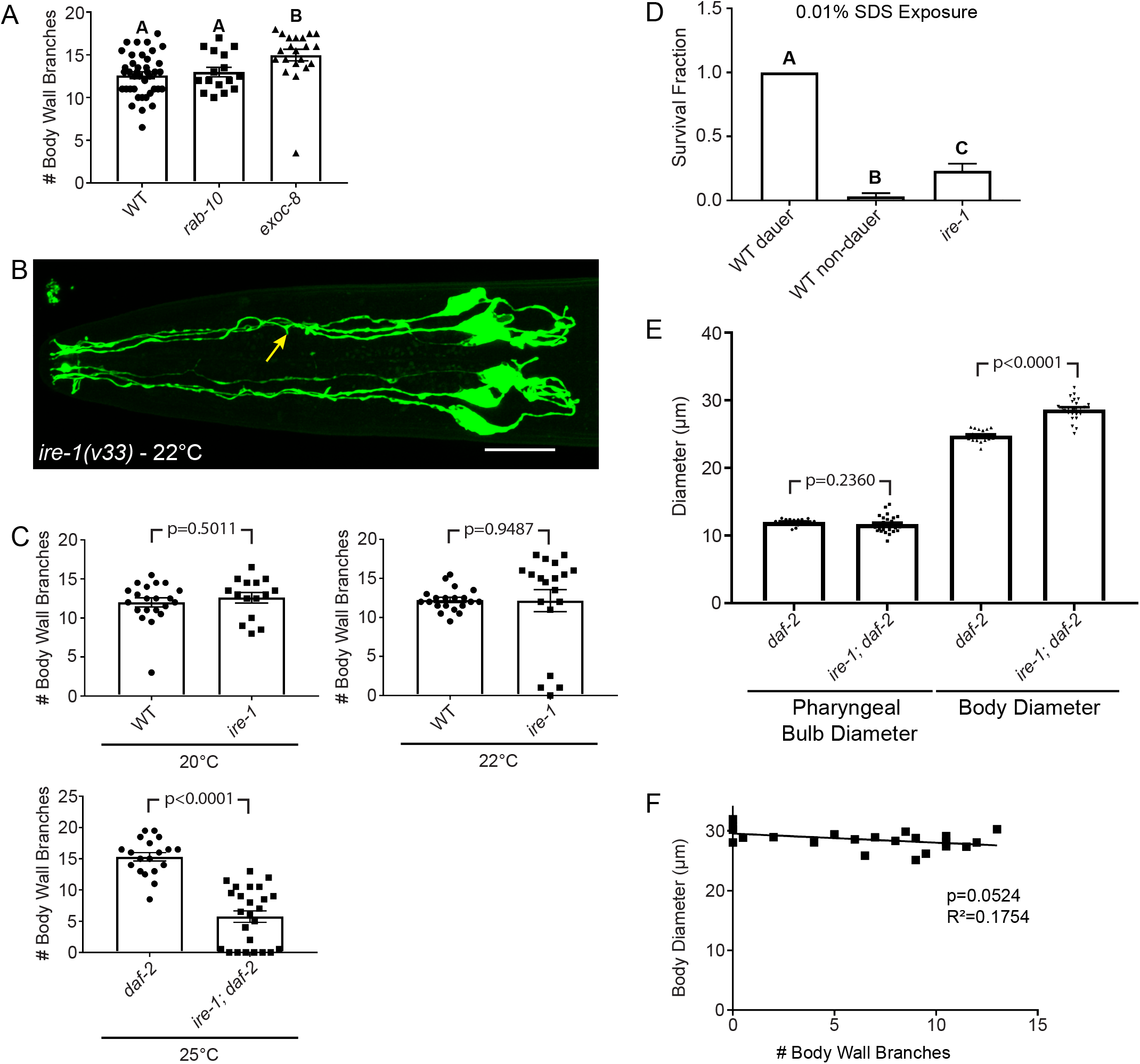
Regulatory mechanisms controlling arborization differ between IL2s and FLP/PVDs. (A) Mutation in *rab-10* does not affect the number of IL2 body wall branches (n=16). The *exoc-8* mutant has a slight increase in the number of body wall branches (n=21, p=0.0039). (B) A Z-projection micrograph dorsal-ventral view of the *ire-1* mutant dauer with truncated secondary branches (yellow arrow). IL2 neurons are visualized by *klp-6p∷gfp.* Scale bar, 10 μm. (C) Quantification of *ire-1* IL2 arbors at multiple temperatures. At 20°C (n=15) and at 22°C, *ire-1* (n=19) mutants show no difference in IL2 branching compared to wild-type (n=20 and n=21, respectively). At 22°C, 21% of the *ire-1* mutants show severe branching defects with fewer than three body wall branches. Due to severe developmental and fertility defects in *ire-1* mutants at 25°C, we were prevented from collecting suffecient dauers at one time point for a successful experiment. To collect data at this time point, we built an *ire-1; daf-2* double mutant that forms dauers constitutively at 25°C. At 25°C, *ire-1; daf-2* dauers (n=25) have decreased branching compared to a *daf-2* control (n=19) (p<0.0001) and 36% of animals have severe branching defects. Animals were considered severely defective if they had fewer than three body wall branches. (D) *ire-1* mutant dauers are defective in SDS survival. WT dauers survive exposure to SDS. *ire-1* survives 0.01% SDS better than the non-dauer WT control (n=60 for each treatment). None of the *ire-1* mutant dauers survived a standard 1% SDS assay (See Fig. S4C) (E) *ire-1* mutants undergo incomplete tissue remodeling during dauer at 25°C. While *ire-1* mutants completely remodel their pharyngeal bulb at 25°C (n=25), the overall body diameter is larger in the *ire-1* mutant than WT (n=19, p<0.0001). (F) Incomplete remodeling of the body wall is weakly correlated with IL2 arborization in *ire-1* mutant dauers. Regression of IL2 branching with body diameter in *ire-1* mutants grown at 25°C (p=0.0524, R^2^=0.1754). For SDS survival assay and IL2 branch count experiments between three groups we used ANOVA followed by Tukey’s test for multiple comparisons to determine significance (p<0.01)(A and D). Different letters above bars indicate statistical significance. For all IL2 branch count experiments between two groups, pharyngeal bulb, and body diameter comparisons, we used t-tests to determine significance (C and E). Error bars are the standard error of the mean.

Similar to the requirement for RAB-10, the unfolded protein response (UPR) sensor IRE-1 is needed for proper localization of DMA-1 and arborization in the PVD neurons (12, 13). Loss of DMA-1 leads to high penetrance defects in PVD branching. Interestingly, we found severe, but low penetrance (21% n=19 defects in IL2 arborization of *ire-1* mutant dauers when grown at 22°C (Fig. 3B). While the total branch number did not differ statistically from wild-type (Fig 3C), we were curious if other factors contributed to the UPR regulation of dauer. To determine if the UPR was activated differently during dauer, we examined mutants of other UPR candidate genes including *skr-5*,*tps-1*,*gst-4*, *F47F2.1*, and *xbp-1* (27–29). However, all other UPR candidates examined were able to form complete IL2 arbors during dauer (Fig. S4B).

During the course of our studies, we observed that populations of *ire-1* mutants crashed if not stored in a temperature-controlled incubator and that there was an apparent correlation with culture viability and fluctuations in room temperature. Indeed, we found that *ire-1* mutants kept at 25°C were sterile with variable developmental defects. We hypothesized that dauer-specific branching in *ire-1* mutants is temperature sensitive. Consistent with this, we found no defects in IL2 arborization at 20°C (Fig. 3C). Unfortunately, the severe defects found in *ire-1* mutants at 25°C precluded us from collecting sufficient dauers for examination of the IL2s. We, therefore, examined the *ire-1* dauers in a *daf-2* constitutive dauer formation genetic background. At 25°C *ire-1; daf-2* mutants have a significant decrease in the number of body wall branches compared with *daf-2* mutants alone (Fig. 3C). These results suggest an interaction between temperature and the UPR in regulating multiple aspects of development.

A common method for isolating dauers from non-dauers is exposure to 1% sodium dodecyl sulfate (SDS). Wild-type dauers can survive exposure to 1% SDS for several hours while it is lethal to non-dauers within minutes (30). During our initial attempts, we were rarely successful at isolating any *ire-1* mutant dauers using SDS at room temperature. We confirmed that *ire-1* mutant dauers were defective for survival to SDS, although could survive higher concentrations of SDS than non-dauer animals (Fig. 3D and S4C). Our SDS results suggest that *ire-1* mutants form only partial dauers. In addition to IL2 arborization, dauers undergo a radial shrinkage of their body and pharynx (30). We found that *daf-2; ire-1* mutant dauers have defects in radial shrinkage at 25°C while pharyngeal remodeling appears normal (Fig. 3E). To determine if the remodeling of different tissues types are correlated, we regressed body diameter against IL2 branching data. We found only weak correlation between branching defects and body diameter (p=0.0524, R^2^=0.1754) (Fig. 3F). These data suggest that *ire-1* acts independently in multiple tissue types to control dauer remodeling.

### DAF-16 is necessary for IL2 arbor formation during dauer, but not FLP arbor formation in adult animals

Our finding that *ire-1* mutants form partial dauers motivated us to examine other regulators of dauer morphogenesis. Previous results showed that loss of the DAF-2 insulin signaling receptor rescues the arborization defects in *ire-1* mutant PVDs (13). This insulin signaling-mediated rescue of the *ire-1* mutant branching phenotype in the PVDs is dependent upon the FOXO transcription factor DAF-16. However, we found that *ire-1; daf-2* double mutants, where DAF-16 is constitutively activated, fail to restore the IL2 arbors to wild-type levels (Fig. 3C). To explore this pathway more directly, we examined IL2 branching in *daf-16* mutants. Loss of DAF-16 causes defective for dauer formation; however, *daf-16* mutants will form partial dauers in a dauer-constitutive (*daf*-c) mutant background (31, 32). *daf-16* partial dauers display some, but not all of the characteristic tissue remodeling found in wild-type dauers. We found that *daf-16; daf-7* partial dauer IL2 arbors were reduced compared to wild-type dauers (Figs. 4A and B). We also found frequent defects in cell body positioning and abnormal axon morphology in *daf-16; daf-7* partial dauers. However, we found no apparent defects in FLP arbor formation in *daf-16; daf-7* adult animals (Fig. S5A) suggesting that DAF-16/FOXO differentially regulates arborization in the IL2s and FLPs.

**Figure 4.**
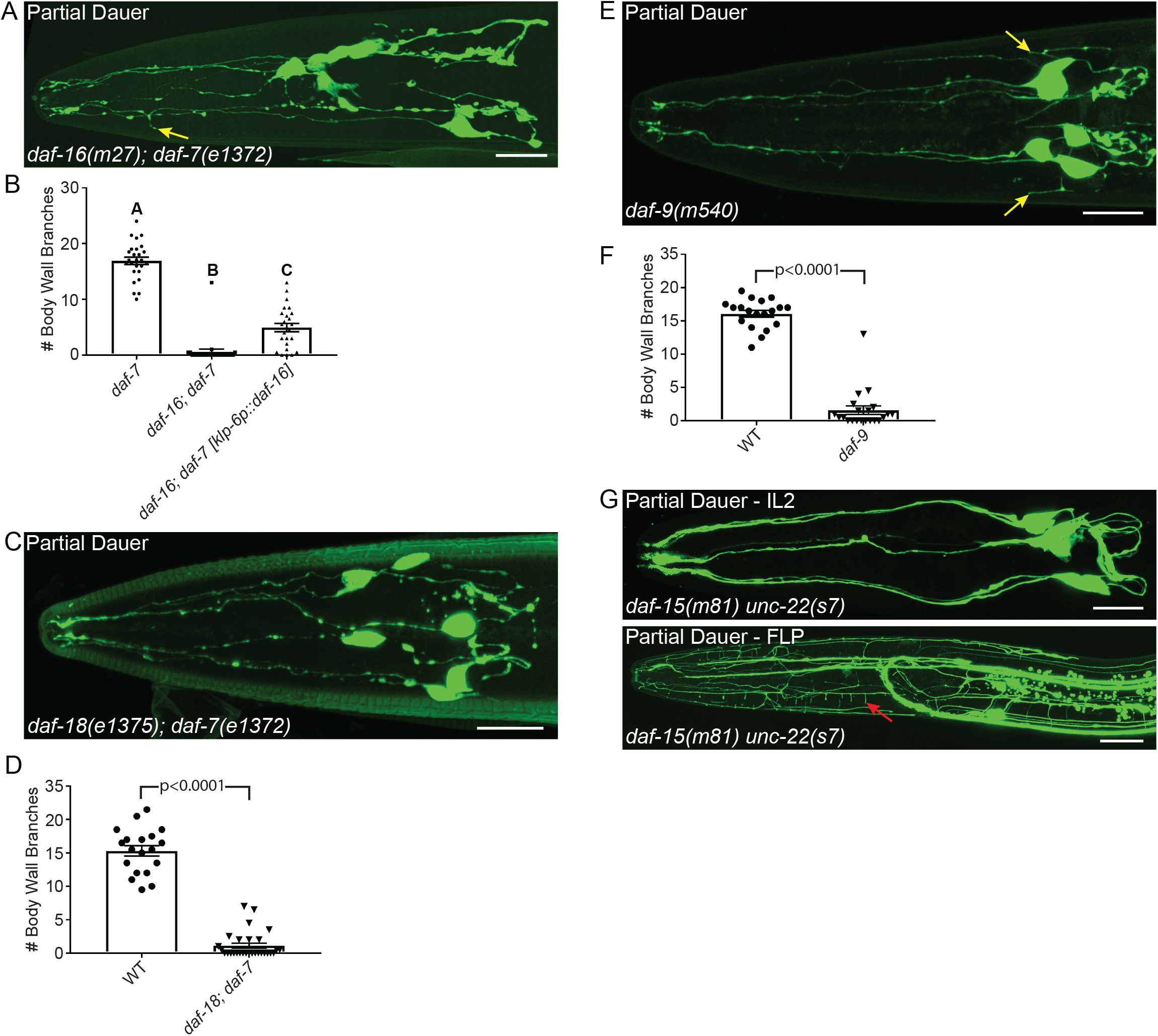
Genes regulating dauer formation are required for IL2 and FLP arborization. (A) Z-projection confocal micrograph of *daf-16(m27); daf-7(e1372)* partial dauers showing few truncated secondary IL2 branches (yellow arrow). (B) Quantification of IL2 body wall branches. The *daf-16(m27); daf-7(e1372)* mutant (n=27) is severely defective in IL2 arbor formation compared to the *daf-7* control (n=26). Expression of *daf-16* from the IL2 neurons with a *klp-6* promotor (n=25) is sufficient to increase arborization. (C) Z-projection confocal micrographs. A dorsal-ventral view of a *daf-18(e1375); daf-7(e1372)* partial dauer animal shows a lack of IL2 body wall branches. (D) Quantification of *daf-18(e1375); daf-7(e1372)* mutant IL2 branching defects. *daf-18(e1375); daf-7(e1372)* partial dauers (n=29) have few branches compared to the *daf-7(e1372)* control (n=19). (E) Z-projection confocal micrograph of IL2 neurons expressing gfp in *daf-9(m540)* partial dauers. *daf-9(m540)* mutants extend several branches directly from the cell body (yellow arrows), but form few higher-order branches. (F) Quantification of the *daf-9(m540)* mutant IL2 defects. *daf-9(m540)* mutants (n=21) have fewer body wall branches compared to the wild type control (n=19). (G) Z-projection confocal micrographs. *daf-15(m81) unc-22(s7)* partial dauers have completely unbranched IL2 neurons (top) and extensive branching of the FLP neuron (bottom), including body wall branches (red arrow). For analyses between two groups, statistical significance was determined with t-tests (D and F). For genetic rescue comparisons, statistical significance was determined with ANOVA followed by Tukey’s test for multiple comparisons (B). Different letters above bars indicate statistical significance (p<0.0001). Error bars are the standard error of the mean. IL2 neurons are visualized by *klp-6p∷gfp* (A,C,E and G) and FLP neurons are visualized by *mec-7p∷gfp* (G). Scale bars,10 μm.

DAF-16 is widely expressed in multiple tissue types (33). Previous work showed that pan-neuronal rescue of DAF-16 is sufficient to rescue all dauer remodeling and formation defects in the *daf-16* mutant (34). To determine if *daf-16* is functioning cell-autonomously, we drove expression in the IL2 neurons alone. Consistent with a cell autonomous role, we found a significant increase in arbor formation following IL2-specific expression of *daf-16* (Fig. 4B). *daf-16* mutants dauers are defective for pharynx remodeling and radial shrinkage (31). We did not observe any change to pharyngeal remodeling or radial shrinkage following IL2-specific rescue (Fig. S5B). Therefore, while DAF-16 expression in the IL2 neurons is sufficient to increase IL2 arborization, it is not capable of inducing system-wide remodeling of dauer tissue. These results suggest that DAF-16 functions cell autonomously to direct IL2 remodeling.

Previous work showed that if *daf-2* mutant adults are transferred to 25°C, they will become reproductively quiescence and exhibit some dauer-like morphology and behaviors (35). We hypothesized that activation of the DAF-16 transcription factor in post-dauer animals would be sufficient to induce IL2 branching. We transferred L4 *daf-2* animals to 25°C and checked for ectopic IL2 growth for three consecutive days. Although we reproduced the reproductive quiescence phenotype, the IL2 dendrites remained unbranched (n=60). Similar to our finding that expression of DMA-1 outside of dauer does not induce IL2 branching, these results suggest a potent inhibition of IL2 branching outside of the dauer stage.

Mutations in several other components of the dauer formation pathway will produce partial dauers either alone or in combination with *daf*-c mutants. The phosphatase and tensin (PTEN) homolog DAF-18 is required for proper dauer formation. Similar to *daf-16, daf-18* mutants form partial dauers in a *daf*-c background (31). We found that *daf-18* is defective in IL2 arbor formation (Fig 4C and D). *daf-9* encodes a cytochrome p450 that synthesizes a lipid hormone which binds to a nuclear hormone receptor to bypass the dauer stage. Animals carrying the *daf-9(m540)* mutation constitutively form dauers when grown on cholesterol-free media (36). We found that *daf-9(m540)* dauers have a reduced number of branches from their primary dendrite, while still exhibiting a multipolar cell body (Fig. 4E and F). *daf-9* mutant adults were wild-type for FLP arborization. Similar to *daf-9*, disruption of the TOR complex associated protein DAF-15 causes arrest of partial dauers. *daf-15* arrested animals have some intermediate dauer-like morphologies (37); however, we found no branching in any *daf-15* partial dauers. To our surprise, the *daf-15* partial dauer FLPs were completely arborized. This was unexpected as the FLPs are typically unbranched throughout the majority of larval development and may suggest that TOR complex 1 regulates the timing and dauer-specific inhibition of FLP arborization (Fig. 4H).

## DISCUSSION

We find that the DMA-1 complex is used to regulate branching in each highly arborized neuron in *C. elegans*. This highlights the versatility of the DMA-1 receptor-ligand complex. DMA-1 is expressed in the IL2s only during dauer. However, we show that the DMA-1 complex is not sufficient to induce branching in the IL2s. Overexpression of DMA-1 in the IL2s during non-dauer stages did not induce branching despite the presence of SAX-7, MNR-1, and LECT-2 (9, 18). Similarly, we found that during dauer the FLPs stop branching despite its expression of DMA-1. Our findings may suggest the presence of inhibitors of IL2 branching outside of dauer and FLP branching during dauer and support the hypothesis that activation of DMA-1 is context-dependent (16). Indeed, our finding of extensive FLP branching in *daf-15* partial dauers suggests that the TORC1 complex is required to inhibit FLP branching during dauer. In *Drosophila*, the TORC2 regulates dendritic tiling (38). In mammals, mTOR localizes to dendrites and is required for local translation (39). Knockdown of mTOR leads to reduced dendrite arborization in cultured hippocampal neurons (40).

Our detailed examination of FLP branching revealed unknown structural differences between it and the well-studied PVD neurons. Both neuron types extend terminal branches along the body wall between the somatic muscle and surrounding hypodermis. Despite the similarity in transcriptional regulation and downstream effectors of branching, the lower-order branches differ substantially between PVD and FLP. The PVD secondary dendrites extend regularly from the laterally positioned primary dendrite, while the FLP secondary dendrites are sparsely distributed and differ between the dorsal and ventral body quadrants. The presence of FLP body wall branches that extend from both the lateral and dorso-ventral midlines appears intermediate between the PVD and IL2 branching patterns. Neurite formation has long been known to be shaped by surrounding tissues. The primary dendrite of the PVDs fasciculates with the relatively sparsely populated lateral nerve containing only 2-3 other neurons (41). In contrast, FLP fasciculates within a sensory neuron fascicle with more than 20 neuronal and glial processes. Also, the hypodermis surrounding the PVDs comprises a single hypodermal syncytium and a line of epithelial seam cells, whereas the area of FLP branching comprises multiple hypodermal and seam cells. These differences in local cues may affect the different architectures seen in each arborizing neuron.

One possible mechanism for the different arborization pattern between the PVD/FLP classes and the IL2 is through the transport of DMA-1. Indeed, we found that, unlike in the PVDs, RAB-10 is not required for IL2 arborization. Similarly, the RAB-10 paralogs RAB-8 and RAB-35 were not needed for branching during dauer. One possible explanation is that the relatively smaller area of the IL2 arbor, compared with the expansive PVD and FLP dendritic trees, does not require a RAB-based membrane transport (Fig S1). Alternatively, an unknown RAB protein may be required specifically in the IL2s.

Interestingly, we found that the inositol requiring enzyme IRE-1, which regulates DMA-1 transport in the PVDs, also plays a substantial role in regulating IL2 arborization under elevated temperature. Studies in plants and fungi demonstrated that IRE-1 homologs are conditionally required under elevated temperatures (42, 43). In the UPR, IRE1 acts as a sensor and activator of the XBP-1 transcription factor. Similar, to findings in the PVDs, we found that *xbp-1* is not required for proper IL2 arborization (12, 13). It is, therefore, likely that the IL2s also use the XBP-1-independent RIDD (regulated Ire1-dependent decay) pathway. Loss of the DAF-2 insulin signaling receptor can suppress the effect *ire-1* mutation in the PVDs (13). However, we found that loss of *daf-2* did not rescue the *ire-1* mutant at elevated temperature. We hypothesize that the normal upregulation of DAF-16 in dauers masks the loss of *ire-1;* however, at higher temperatures and corresponding increased levels of cellular stress, the IRE-1 mechanism becomes necessary for IL2 branching.

Our examination of *ire-1* mutants suggest that ER-related homeostasis regulates several aspects of dauer remodeling. While our statistical analysis suggests independence between different tissue remodeling events, future work will be needed to determine if dauer remodeling in other tissues is acting through the canonical UPR or XBP-1 independent mechanisms. The UPR was previously implicated in dauer formation in the *daf-28 daf*-c mutant background (44). Similarly, the UPR regulates various aspects of remodeling in mammalian tissues (45, 46).

Dauer formation comprises remodeling of neurons, muscle, and epithelial tissue (47). In wild-type animals these various tissues must coordinate their remodeling to produce a complete dauer; a concept in evolution known as phenotypic integration. While the mechanisms controlling the decision to enter dauer have been extensively studied, the mechanisms regulating individual dauer remodeling events and the coordination of these events are less well studied. We found that several mutants with known defects in dauer morphology also have defects in IL2 arborization. The FOXO transcription factor homolog DAF-16 is translocated to the nucleus under conditions of environmental stress. *daf-16* partial dauers undergo incomplete remodeling of epithelial and alimentary tissues (31). DAF-16 is required for proper axon outgrowth in the *C. elegans* AIY interneurons and mammalian cerebellar granule neurons (48). We add to this by demonstrating a lack of IL2 branching in *daf-16* partial dauers. Similar to its role in AIY outgrowth, we find that DAF-16 is acting cell autonomously.

We also found defects in *daf-15* and *daf-18* mutants. *daf-18* encodes the Phosphatase and Tensin (PTEN) homolog and, in combination with *daf-c* mutants, produces partial dauers with an unremodeled pharynx that continues to pump (31). PTEN plays a major role in neuron development. Loss of PTEN in mice leads to hypertrophic arborization in the dentate gyrus (49). Inhibition of neurite outgrowth by PTEN in mammalian systems acts through mTOR activation (50). The TOR pathway *daf-15* mutation leads to irreversible dauer formation with very few dauer characteristics. We find that *daf-15* mutant partial dauers lack all branching. Given the lack of dauer-specific features, we question whether these were true dauers. A recent publication showing reduction of dauer-entry expression of the *ets-10p∷gfp* transgene in a *daf-15* background (51). Additional experiments will be required to understand the possible interactions of DAF-18 and DAF-15 in IL2 arborization.

## MATERIALS AND METHODS

### Maintenance of *C. elegans* strains

*C. elegans* were cultured and genetic crosses were performed on nematode growth media (NGM) with *E. coli* OP50 bacteria as a food source according to standard protocol (4). All strains were maintained at 22°C unless otherwise noted. All mutants were backcrossed at least twice to wild-type. *dma-1(cbc1)* was generated using a standard EMS mutagenesis protocol (52). A complete list of strains used in this study can be found in Table S1. Dauers were produced using several methods including through exposure to crude pheromone, selection from starved and overcrowded culture plates, and by using a dauer formation constitutive mutant background. Dauers were tentatively identified under the dissecting microscope by their relatively thin body, lack of pharyngeal pumping and head foraging movement. Following examination under the compound microscope, the animals were confirmed to be dauer if they had an obvious buccal plug and lateral alae.

### Plasmid constructs and generation of transgenic lines

Plasmids were built in the Schroeder Lab using Gibson Assembly (53). Rescuing constructs were injected following standard microinjection technique at 5 ng/μl together with coel∷RFP at 50 ng/μl and pBluescript to a final concentration of 100 ng/μl (54). At least two independent stable lines were isolated and analyzed for each construct. In all cases, the lines gave similar results and, therefore, results from one line are presented. Successful rescue was determined with ANOVA followed by Tukey’s multiple comparison. A complete list of plasmids, and assembly primers can be found in Table S2.

### Genetic Mapping

Newly generated mutant strains were outcrossed twice and assigned to a linkage group based on two-factor mapping with EG1020 *bli-6(sc16) IV; dpy-11(e224) V; lon-2(e678) X* and EG1000 *dpy-5(e61)I;rol-6(e187)II;lon-1(e1820)III*. The new allele *dma-1* was confirmed through complementation testing performed on dauer pheromone-containing media with *dma-1(tm5159)I*. The *dma-1(cbc1)* lesion was identified with Sanger Sequencing.

### Gene Schematics

Gene schematics were constructed using wormweb.org. Functional domains for *dma-1* were determined using LRRfinder.com.

### FLP developmental timecourse

We used *muIs32[mec-7p∷gfp]* to visualize FLP cell morphology. To generate synchronous cultures, eggs were collected from gravid hermaphrodites by bleaching with 5% bleach and 5N NaOH in M9 buffer (55). Eggs were allowed to hatch overnight in M9, generating a population of L1 arrested larva. Arrested L1 larvae were then transferred to NGM plates seeded with OP50 *E. coli*. The L1 larvae were allowed to develop at 22°C until the desired stage for imaging. We evaluated the worms after 1, 14, 22, 30, and 40 hours on food to determine FLP morphology at the L1, L2, L3, L4 and young adult stages, respectively. Dauers were selected based on dauer morphology and behavior from recently food-depleted plates. Representative images for each stage were acquired using confocal microscopy due to the complexity of the FLP arbor. In some cases multiple z-stacks were combined using Image J to obtain a composite image of the full arbor. Concurrently a sample size of 20 animals were observed under a compound microscope at each time point to confirm the consistency of our observations.

### IL2 body wall branch counts

The Number of branches originating from either the dorsal or ventral midlines were counted to obtain a body wall branch number. The branch counts were averaged for all cells quantified within an individual. Averages were then compared using either a t-test or ANOVA followed by Tukey’s test for multiple comparisons to determine statistical significance.

### Microscopy

Animals were anesthetized with 100 mM levamisole and mounted on 10% agarose pads containing 20 mM levamisole for sustained immobilization during imaging. Datasets for quantification were gathered with a Zeiss AxioImager microscope equipped with DIC and fluorescence optics. High-resolution representative images were acquired with a Zeiss LSM 880 confocal microscope with Airyscan postprocessing. Body diameter was measured near the terminal pharyngeal bulb. Pharyngeal bulb diameter was measured at the widest point of the bulb. Measurement data was compared to a control dauer dataset using a t-test.

### Sodium dodecylsulfate sensitivity assays

Sodium dodecylsulfate (SDS) sensitivity assays were performed in 12-well petri plates containing M9 buffer and the specified concentration of SDS. Animals were incubated at room temperature with gentle shaking for 30 minutes and scored for survival by looking for movement following touch with a platinum wire. Each treatment was tested in triplicate. Each replicate contained a wild-type dauer control and a wild-type L3 non-dauer control. Significant difference in survival rates were determined with ANOVA followed by Tukey’s test for multiple comparisons.

### Temperature shift assay

*daf-2(e1370);myIs14 [klp-6p∷gfp]* one-day-old adults were transferred from 15°C to 25°C. Twenty animals were examined for IL2 morphology every 24 hours for three consecutive days.

## ACKNOWLEDGEMENTS

We thank the Hobert, Shen, Buelow, and Domier labs for sharing strains, plasmids, and reagents. Some strains were provided by the CGC, which is funded by the NIH Office of Research Infrastructure Programs (P40 OD010440). Funding in the Schroeder lab was provided by NIH-NIGMS R01GM111566 to NES. BR and CMS were supported by an NSF Research Experience for Undergraduate Site award to NES (1559908/1559929).

## SUPPLEMENTARY FIGURES

**Figure S1. Arborization of different neuronal subtypes during *C. elegans* development.** (A) Under favorable conditions, *C. elegans* develops from an egg, through four larval stages and into a mature adult. When grown under conditions of low food and high population density, *C. elegans* larvae enter into an alternative developmental stage called dauer. When dauers are returned to food they exit dauer and continue development into reproductive adults. There are slight morphological differences between post-dauer adults and those that have not gone through dauer. (B) There are three classes of highly arborized neurons in *C. elegans*: IL2, FLP, and PVD. The IL2s (red) arborize strictly during the dauer stage where their arbors are spatially restricted to the head of the nematode. The FLPs (blue) and PVDs (purple) form arbors during late L4 and together cover the head and body, respectively. Life cycle modified from Androwski et al. 2017 (1).

**Figure S2. DMA-1 binding partners are required for IL2 arborization.** (A) Z-projection confocal micrographs of dauers expressing gfp in the IL2s. Wild-type dauers have extensive IL2 arbors. *dma-1(tm5159)*, *mnr-1(wy758)*, and *lect-2(ok2617)* mutants are severely defective in IL2 branching. (B) Expression of *mnr-1* in the hypodermis using the *dpy-7* promoter rescues the *mnr-1* mutant branching phenotype. Quantification of the number of IL2 branches along the body wall during dauer in wild-type (n=21), *mnr-1*(n=20), and *mnr-1; dpy-7p∷mnr-1* (n=16). (C) Expression of *lect-2* in the hypodermis using the *myo-3* promoter rescues the *lect-2* mutant branching phenotype. Quantification of the number of IL2 branches along the body wall during dauer in wild-type (n=19), *lect-2* (n=19), and *lect-2; myo-3p∷lect-2* (n=16). For all rescue experiments an ANOVA followed by Tukey’s test for multiple comparisons was used to determine significance. Different letters above bars indicate statistical significance (p<0.0001). IL2 neurons are visualized by a fluorescent reporter, *klp-6p∷gfp* (A). Error bars are the standard error of the mean. Scale bars, 10 μm.

**Figure S3. The FLP dendrite bifurcates early in development. Z-projection confocal micrographs of FLP neurons throughout larval development.** (A) Dorsal-ventral view of the FLP dendrite in L1. Each FLP primary dendrite extends from the cell body to anterior of the metacorpus where it branches to send two processes to the nose. (B-C) Lateral left view of the L2 and L3 FLP dendrite. Little additional branching occurs during the L2 and L3 larval stages. (D) The FLPs arborize rapidly during L4. Arrows indicate part of the FLP dendrite. Asterisks indicate FLP cell bodies. The FLPs are labeled with *mec-7p∷gfp*, which also labels the ALM and AVM touch receptor neurons. In addition to the FLPs, the unbranched ALM and AVM head neurons also labeled by this reporter. Scale bars, 10 μm.

**Figure S4. Analysis of RAB-10 paralogs and unfolded protein response candidate genes and their effect on dauer morphology.** (A) Quantification of IL2 arborization in *rab-10* paralogs. Mutations in *rab-8* (n=19) or *rab-35* (n=21) do not affect IL2 branch number compared to wild-type (n=17 and n=22, respectively). (B) Quantification of IL2 arborization in unfolded protein response mutants. *skr-5* (n=21), *F47F2.1* (n=21), and *tps-1* (n=21) do not differ from wild-type in the number of body wall branches (n=18), while *xbp-1* (n=19) has a slight increase in branch number (p=0.0411). (C) Quantification of *ire-1* SDS survival. At SDS concentrations of 0.1%, 0.2%, 0.5%, and 1%, *ire-1* dauers fail to survive better than the non-dauer controls (n=60 for all treatments, p<0.0001). For IL2 body wall branch count experiments comparing two strains, we used t-tests to determine statistical significance (A and B). For SDS survival assays and IL2 branch counts between more than two groups we used ANOVA followed by Tukey’s test for multiple comparisons to determine significance (B and C). Error bars are the standard error of the mean. Different letters above bars indicate statistical significance.

**Figure S5. The role of dauer formation genes in FLP arborization and dauer-specific tissue remodeling.** (A) Z-projection confocal micrographs of *daf-16(m27); daf-7(e1372)* adults. Lateral view of the FLP neurons in an adult shows a fully arborized FLP dendrite. (B) Quantification of pharyngeal bulb and body diameter in partial dauers. The *daf-16(m27); daf-7(e1372)* mutant (n=27) is defective in pharyngeal bulb remodeling and constriction of body diameter compared to the *daf-7* control (n=26). Expression of *daf-16* from the IL2 neurons with a *klp-6* promotor (n=25) is unable to rescue defective remodeling of the pharyngeal bulb or radial constriction of the body during dauer. Statistical significance was determined with ANOVA (B). Different letters above bars indicate statistical significance (p<0.0001). (C) Z-projection confocal micrograph lateral view of a *daf-18(e1375); daf-7(e1372)* adult with extensive FLP arbor. (D) Z-projection confocal micrograph of FLP neurons in *daf-9(m540)* mutant adults. Error bars are the standard error of the mean. FLP neurons are visualized by *mec-7p∷gfp* (A, C and D). Scale bars,10 μm.

## REFERENCES

1. Popov VI, Bocharova LS (1992) Hibernation-induced structural changes in synaptic contacts between mossy fibres and hippocampal pyramidal neurons. Neuroscience 48(1):53–62.

2. Woolley CS, Gould E, McEwen BS (1990) Exposure to excess glucocorticoids alters dendritic morphology of adult hippocampal pyramidal neurons. Brain Res 531(1–2):225–231.

3. Schroeder NE, et al. (2013) Dauer-specific dendrite arborization in C. elegans is regulated by KPC-1/furin. Curr Biol 23(16):1527–1535.

4. Brenner S (1974) The genetics of Caenorhabditis elegans. Genetics 77(1):71–94.

5. Smith CJ, et al. (2010) Time-lapse imaging and cell-specific expression profiling reveal dynamic branching and molecular determinants of a multi-dendritic nociceptor in C. elegans. Dev Biol 345(1):18–33.

6. Golden JW, Riddle DL (1984) The Caenorhabditis elegans dauer larva: Developmental effects of pheromone, food, and temperature. Dev Biol 102(2):368–378.

7. Ward S, Thomson N, White JG, Brenner S (1975) Electron microscopical reconstruction of the anterior sensory anatomy of the nematode Caenorhabditis elegans. J Comp Neurol 160(3):313–337.

8. Dong X, et al. (2016) Precise regulation of the guidance receptor DMA-1 by KPC-1/furin instructs dendritic branching decisions. Elife 5(MARCH2016). doi:10.7554/eLife.11008.

9. Salzberg Y, et al. (2013) Skin-derived cues control arborization of sensory dendrites in Caenorhabditis elegans. Cell 155:308–320.

10. Dong X, Liu OW, Howell AS, Shen K (2013) An extracellular adhesion molecule complex patterns dendritic branching and morphogenesis. Cell 155(2):296–307.

11. Zou W, et al. (2016) A multi-protein receptor-ligand complex underlies combinatorial dendrite guidance choices in C. elegans. Elife 5(OCTOBER2016):308–317.

12. Wei X, et al. (2015) The unfolded protein response is required for dendrite morphogenesis. Elife 4:e06963.

13. Salzberg Y, et al. (2017) Reduced insulin/insulin-like growth factor receptor signaling mitigates defective dendrite morphogenesis in mutants of the ER stress sensor IRE-1. PLoS Genet 13(1):e1006579.

14. Zou W, et al. (2015) RAB-10-dependent membrane transport is required for dendrite arborization. PLoS Genet 11(9):e1005484.

15. Taylor CA, Yan J, Howell AS, Dong X, Shen K (2015) RAB-10 regulates dendritic branching by balancing dendritic transport. PLoS Genet 11(12):e1005695.

16. Liu OW, Shen K (2012) The transmembrane LRR protein DMA-1 promotes dendrite branching and growth in C. elegans. Nat Neurosci 15(1):57–63.

17. Zou W, et al. (2018) A dendritic guidance receptor complex brings together distinct actin regulators to drive efficient F-actin assembly and branching. Dev Cell 45(3):362–375.e3.

18. Díaz-Balzac CA, et al. (2016) Muscle- and skin-derived cues jointly orchestrate patterning of somatosensory dendrites. doi:10.1016/j.cub.2016.07.008.

19. Sarov M, et al. (2012) A genome-scale resource for in vivo tag-based protein function exploration in C. elegans. Cell 150(4):855–866.

20. Chattarji S, Tomar A, Suvrathan A, Ghosh S, Mostafizur Rahman M (2015) Neighborhood matters: divergent patterns of stress-induced plasticity across the brain. Nat Publ Gr 18. doi:10.1038/nn.4115.

21. Oren-Suissa M, Hall DH, Treinin M, Shemer G, Podbilewicz B (2010) The fusogen EFF-1 controls sculpting of mechanosensory dendrites. Science (80-) 328(5983):1285–1288.

22. Albeg A, et al. (2011) C. elegans multi-dendritic sensory neurons: Morphology and function. Mol Cell Neurosci 46(1):308–317.

23. Albert PS, Riddle DL (1983) Developmental alterations in sensory neuroanatomy of the Caenorhabditis elegans dauer larva. J Comp Neurol 219(4):461–81.

24. Procko C, Lu Y, Shaham S (2011) Glia delimit shape changes of sensory neuron receptive endings in C. elegans. Development 138(7):1371–1381.

25. Richardson CE, Yee CS, Shen K (2019) A hormone receptor pathway cell-autonomously delays neuron morphological aging by suppressing endocytosis. PLOS Biol:1–20.

26. Stenmark H, Olkkonen VM (1997) The Rab GTPases family. Genome Biol 2001 25 176(5):1–85.

27. Safra M, et al. (2014) The FOXO transcription factor DAF-16 bypasses ire-1 requirement to promote endoplasmic reticulum homeostasis. Cell Metab 20(5):870–881.

28. Shen X, et al. (2001) Complementary signaling pathways regulate the unfolded protein response and are required for C. elegans develoment. Cell 107:893–903.

29. Tawe WN, Eschbach ML, Walter RD, Henkle-Dührsen K (1998) Identification of stress-responsive genes in Caenorhabditis elegans using RT-PCR differential display. Nucleic Acids Res 26(7):1621–1627.

30. Cassada RC, Russell RL (1975) The dauerlarva, a post-embryonic developmental variant of the nematode Caenorhabditis elegans. Dev Biol 46(2):326–342.

31. Vowels JJ, Thomas JH (1992) Genetic analysis of chemosensory control of dauer formation in Caenorhabditis elegans. Genetics 130(1):105–123.

32. Gottlieb S, Ruvkun G (1994) daf-2, daf-16 and daf-23: Genetically interacting genes controlling dauer formation in Caenorhabditis elegans. Genetics 137(1):107–120.

33. Ogg S, et al. (1997) The Fork head transcription factor DAF-16 transduces insulin-like metabolic and longevity signals in C. elegans. Nature 389(6654):994–999.

34. Libina N, Berman JR, Kenyon C (2003) Tissue-specific activities of C. elegans DAF-16 in the regulation of lifespan. Cell 115(4):489–502.

35. Gems D, et al. (1998) Two pleiotropic classes of daf-2 mutation affect larval arrest, adult behavior, reproduction and longevity in Caenorhabditis elegans. Genetics 150(1):129–155.

36. Jia K, Albert PS, Riddle DL (2002) A Caenorhabditis elegans type I TGF beta receptor can function in the absence of type II kinase to promote larval development. Development 129:221–231.

37. Albert PS, Riddle DL (1988) Mutants of Caenorhabditis elegans that form dauer-like larvae. Dev Biol 126(2):270–293.

38. Koike-Kumagai M, Yasunaga K, Morikawa R, Kanamori T, Emoto K (2009) The target of rapamycin complex 2 controls dendritic tiling of Drosophila sensory neurons through the Tricornered kinase signalling pathway. EMBO J 28(24):3879–92.

39. Takei N, et al. (2004) Brain-derived neurotrophic factor induces mammalian target of rapamycin-dependent local activation of translation machinery and protein synthesis in neuronal dendrites. J Neurosci 24(44):9760–9769.

40. Jaworski J, Spangler S, Seeburg DP, Hoogenraad CC, Sheng M (2005) Control of dendritic arborization by the phosphoinositide-3′-kinase-Akt-mammalian target of rapamycin pathway. J Neurosci 25(49):11300–11312.

41. White JG, Southgate E, Thomson JN, Brenner S (1986) The structure of the nervous system of the nematode Caenorhabditis elegans. Philos Trans R Soc London 314(1165):1–340.

42. Deng Y, et al. (2016) IRE1, a component of the unfolded protein response signaling pathway, protects pollen development in Arabidopsis from heat stress. Plant J 88(2):193–204.

43. Cheon SA, et al. (2011) Unique evolution of the UPR pathway with a novel bZIP transcription factor, HxL1, for controlling pathogenicity of cryptococcus neoformans. PLoS Pathog 7(8). doi:10.1371/journal.ppat.1002177.

44. Kulalert W, Kim DH (2013) The unfolded protein response in a pair of sensory neurons promotes entry of C. elegans into dauer diapause. Curr Biol 23(24):2540–2545.

45. Afroze D, Kumar A (2019) ER stress in skeletal muscle remodeling and myopathies. FEBS J 286(2):379–398.

46. Baek J-H, et al. (2019) GRP78 level is altered in the brain, but not in plasma or cerebrospinal fluid in Parkinson’s Disease patients. Front Neurosci 13(July):1–12.

47. Androwski RJ, Flatt KM, Schroeder NE (2017) Phenotypic plasticity and remodeling in the stress-induced Caenorhabditis elegans dauer. Wiley Interdiscip Rev Dev Biol 6(5):e278.

48. Christensen R, de la Torre-Ubieta L, Bonni A, Colon-Ramos DA (2011) A conserved PTEN/FOXO pathway regulates neuronal morphology during C. elegans development. Development 138(23):5257–5267.

49. Kwon CH, et al. (2006) Pten regulates neuronal arborization and social interaction in mice. Neuron 50(3):377–388.

50. Liu K, et al. (2010) PTEN deletion enhances the regenerative ability of adult corticospinal neurons. Nat Neurosci 13(9):1075–1081.

51. Shih P-YY, Lee JS, Sternberg PW (2019) Genetic markers enable the verification and manipulation of the dauer entry decision. Dev Biol. doi:10.1016/j.ydbio.2019.06.009.

52. Anderson P (1995) Mutagenesis. Methods Cell Biol. 48:31–58.

53. Gibson DG, et al. (2009) Enzymatic assembly of DNA molecules up to several hundred kilobases. Nat Methods 6(5):343–345.

54. Mello CC, Kramer JM, Stinchcomb D, Ambros V (1991) Efficient gene transfer in C.elegans: extrachromosomal maintenance and integration of transforming sequences. EMBO J 10(12):3959–3970.

55. Sulston JE, Hodgkin JG (1988) The nematode Caenorhabditis elegans. Science (80-) 240(4858):1448–1453.

