## Supplementary figures and images for "Stress-induced dendritic branching in *C. elegans* requires both common arborization effectors and stress-responsive molecular pathways"

### Supplemental Figure 1

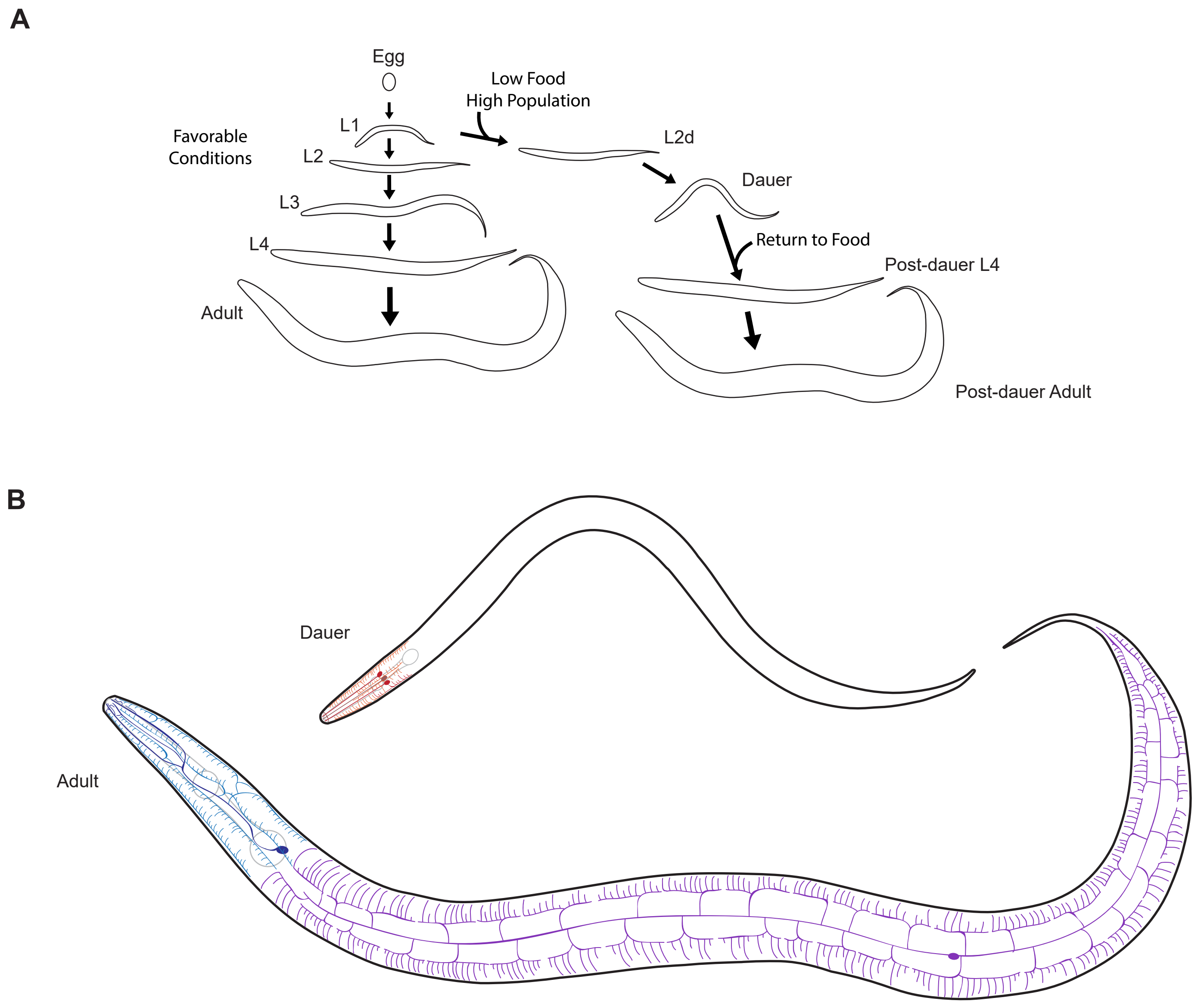

### Supplemental Figure 2

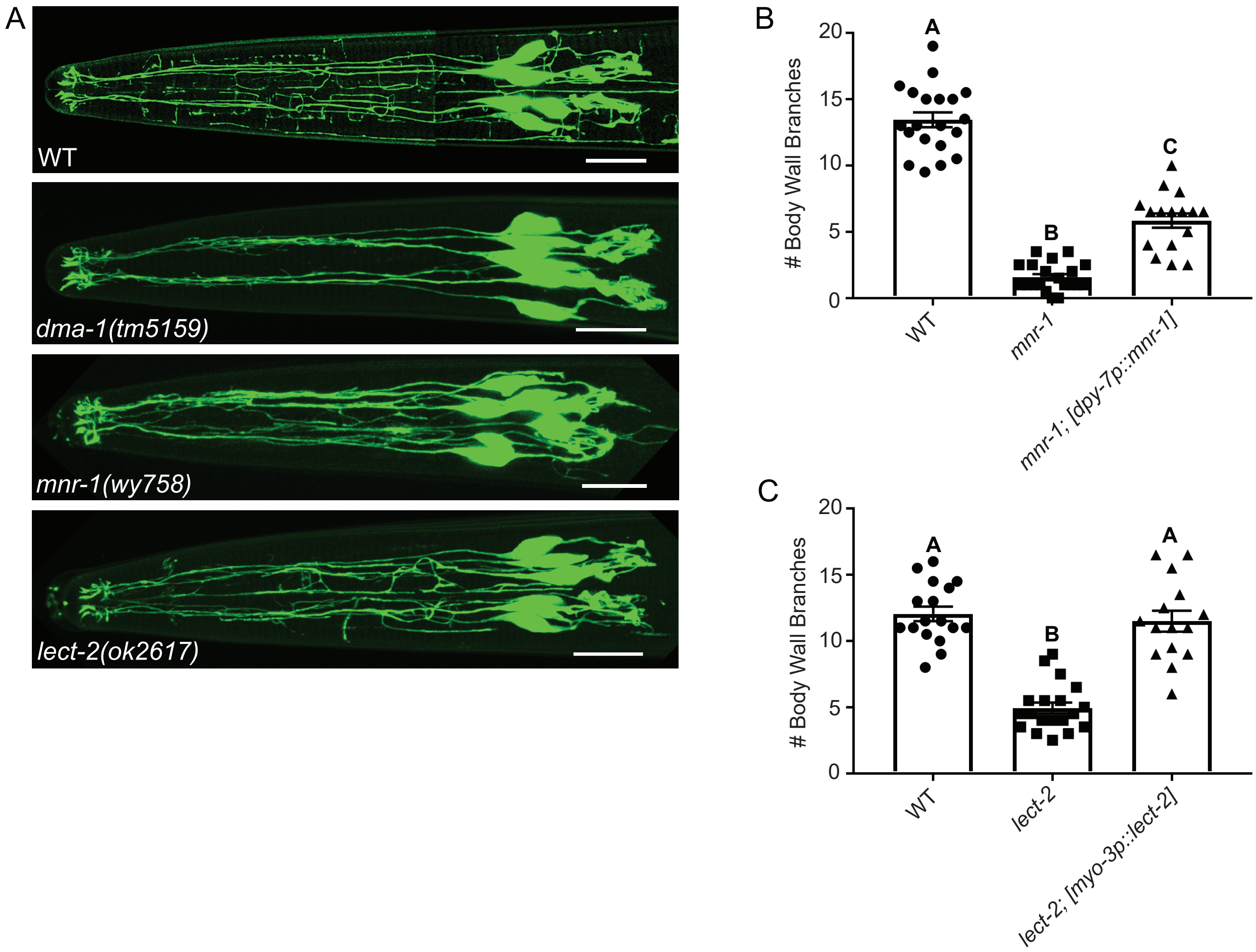

### Supplemental Figure 3

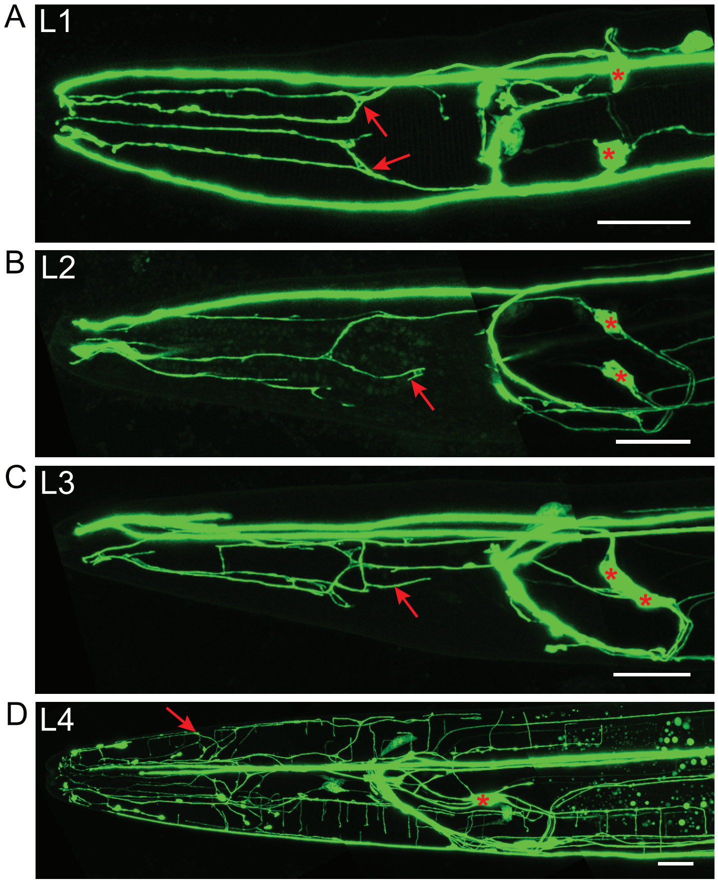

### Supplemental Figure 4

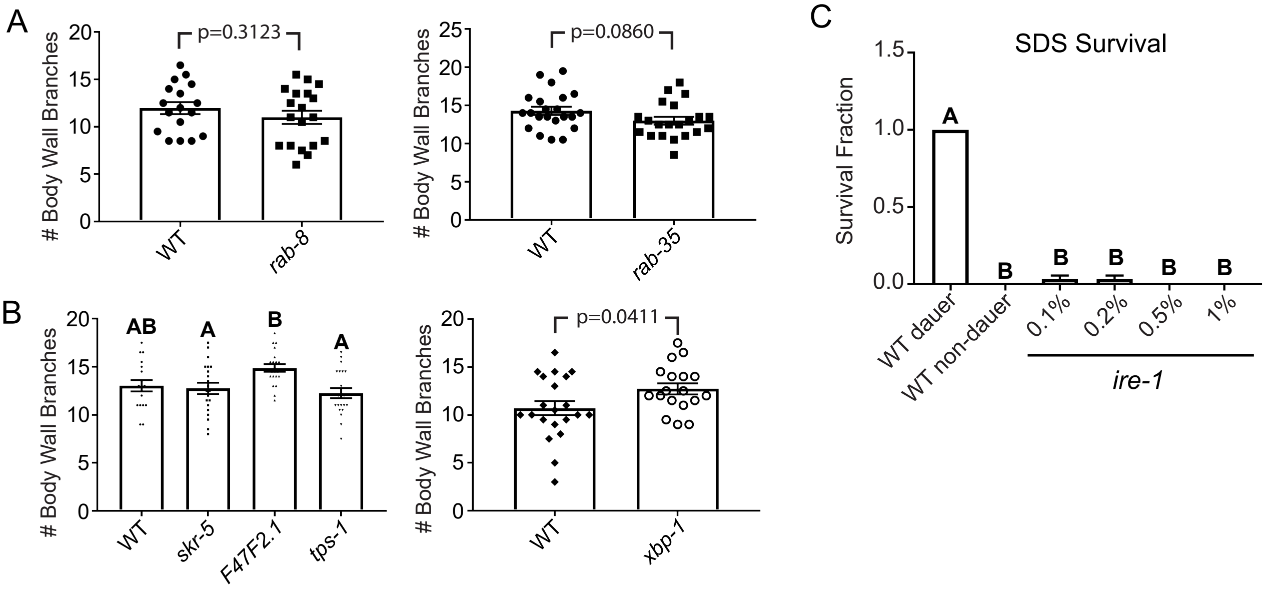

### Supplemental Figure 5

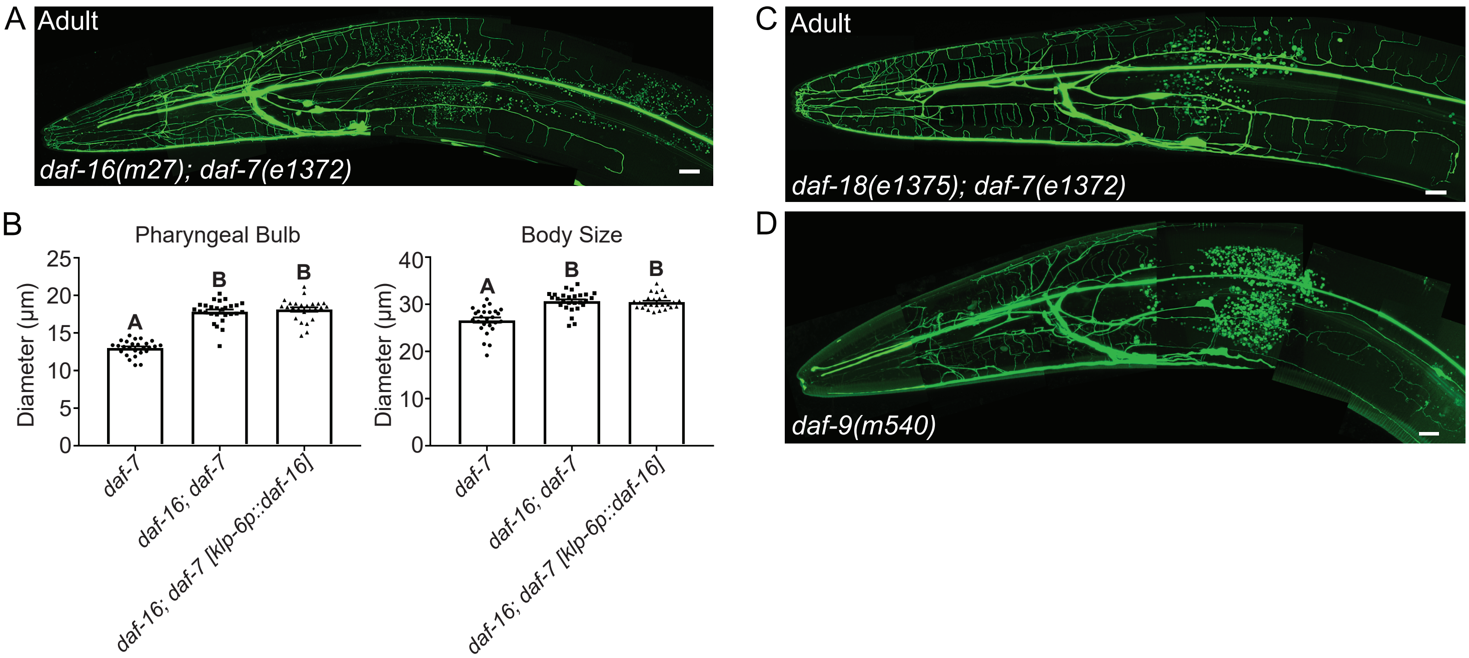
